# Bridging evolutionary game theory and metabolic models for predicting microbial metabolic interactions

**DOI:** 10.1101/623173

**Authors:** Jingyi Cai, Tianwei Tan, Siu Hung Joshua Chan

## Abstract

Microbial metabolic interactions impact ecosystems, human health and biotechnological processes profoundly. However, their determination remains elusive, invoking an urgent need for predictive models that seamlessly integrate metabolic details with ecological and evolutionary principles which shape the interactions within microbial communities. Inspired by the evolutionary game theory, we formulated a bi-level optimization framework termed NECom for the prediction of Nash equilibria of microbial community metabolic models with significantly enhanced accuracy. NECom is free of a long hidden ‘forced altruism’ setup in previous static algorithm while allowing for ‘sensing and responding’ between microbial members that is missing in dynamic methods. We successfully predicted several classical games in the context of metabolic interactions that were falsely or incompletely predicted by existing methods, including prisoner’s dilemma, snowdrift game and mutualism. The results provided insights into why mutualism is favorable despite seemingly costly cross-feeding metabolites, and demonstrated the potential to predict heterogeneous phenotypes among the same species. NECom was then applied to a reported algae-yeast co-culture system that shares typical cross-feeding features of lichen, a model system of mutualism. More than 1200 growth conditions were simulated, of which 488 conditions correspond to 3221 experimental data points. Without fitting any ad-hoc parameters, an overall 63.5% and 81.7% reduction in root-mean-square error in predicted growth rates for the two species respectively was achieved when compared with the standard flux balance analysis. The simulation results further show that growth-limiting crossfeeding metabolites can be pinpointed by shadow price analysis to explain the predicted frequency-dependent growth pattern, offering insights into how stabilizing microbial interactions control microbial populations.

## INTRODUCTION

In nature, microorganisms seldom exist in isolation. Instead, they form communities governed by different types of interactions, which play essential roles in adaptation to environments (1-4) and evolution of species (5-9). Among microbial interactions, exchange of metabolites is especially important and arguably primarily responsible for the fact that more than 99% of bacterial species are not cultivable (4, 10, 11). Understanding metabolic interactions is a fundamental task in microbiome sciences. Despite continual progresses in determining metabolite exchanges using experimental approach such as spatially separated apparatus designs (12, 13), isotope probing (14, 15) and tracing (16) techniques, 16s RNA (17) analysis and metabolome analysis (18), we still need governing principles to predict and understand these metabolic interactions.

Benefiting from the advancement of genome-scale metabolic models (19) and constraint based modeling techniques, modeling of microbial communities has been extended to the level of whole-cell metabolism. Since the first published work on constraint-based modeling of microbial communities more than a decade ago (20), new algorithms addressing various challenges have been proposed. Joint flux balance analysis (Joint-FBA) was first introduced to model a microbial community as a ‘super’ organism containing the compartments for each organism and an additional compartment for inter-cellular metabolite exchange (20, 21). The total community biomass was maximized to predict metabolism. Methods based on Joint-FBA for analyzing host-microbe and microbe-microbe interactions were widely applied to the gut microbiome (22-25). Later a dynamic simulation framework called dynamic multi-species metabolic modeling (DMMM) and other similar methods that iteratively perform FBA for each microbial species at each time step has been developed (26-28). Succeeding efforts include regularization to ensure a well-conditioned dynamical system (29) and incorporation of spatiotemporal elements (30-34). However, in most dynamic methods, the metabolic phenotype of an individual microbe is determined only by the metabolites present in the environment (by finding a particular FBA solution). Unlike Joint-FBA, there is no way for a microbe to directly ‘sense and respond’ to other microbes. These methods might therefore overlook some potential metabolic interactions favored by evolution. In order to reconcile the assumptions of fitness optimization for individuals vs. the entire community, OptCom (35) and its dynamic version d-OptCom (36), solve bilevel optimization problems to optimize the individual fitness in the inner problem, while optimizing the community fitness in the outer problem. To address the effect of viable abundance on the inter-cellular fluxes at population steady state, community FBA (cFBA) (37) generalized joint-FBA by adding parameterized abundances as weights to exchange fluxes. SteadyCom (38) as a reformulation of cFBA enabled efficient computation for flux variability analysis and other constraint based modeling techniques for communities with a large number of organisms. Recently, there have been efforts to simulate communities with a large number of species (39, 40) or handle a large quantity of cases (41-43). Joint-FBA and its linear derivatives are usually employed to avoid expensive computation. However, some constraints and/or the community objective function in joint-FBA and other static FBA-based algorithms force individual organisms to produce a certain amount of metabolites for other members prior to optimizing its own fitness whenever doing so maximizes the community-level objective function value. Here we term this ‘forced altruism’. We believe that forced altruism is potentially applicable to microbes that are physically connected, e.g., by nanotubes (44, 45) or to endosymbiosis, since inter-cellular exchange under these situations are likely governed by concentration gradients or regulation by the host, respectively. In general for microbes in a co-culture separate from each other by cell envelopes and not controlled by higher-level regulatory mechanisms (e.g., host-regulation, quorum sensing, etc.), however, the applicability of ‘forced altruism’ constraints should be questioned.

From evolutionary game theory, Nash equilibria (NE) and evolutionary stable strategies (ESSes) which are a subset of NE are more informative targets to identify for studying the community interactions (46). A microbial community is in a NE if the strategy of each microbe is maximizing its fitness function in the given environment and an ESS is a NE not invadable by other strategies (46). From the previous discussion, while joint-FBA and other static methods do not satisfy the condition of NE, dynamic methods simulating over a long time until equilibrium could reach a particular NE but may miss alternative strategies that are more evolutionarily favorable. In community metabolic networks, without introducing ad-hoc parameters, each microbe is represented as the sum of its constituent macromolecules synthesized from the network and the synthesis rate is usually used to represent fitness. The available strategies for each player are any possible flux distributions (the entire set of reaction fluxes) satisfying the mass balance principle, reaction directionality indicated by thermodynamics, and constrained by substrate availability, which depends not only on the nutrients in the environment but also the strategies of other players in terms of the cross-feeding metabolites they may produce (47). The metabolic details, continuous flux space and interdependence of available strategies between players characterize a unique class of microbial games in community metabolic networks (see Figure 1) that require novel game-theoretical methods. In this paper we introduce a bi-level mixed integer optimization framework free of forced altruism termed NECom to identify various NE. We demonstrate the ability of the proposed algorithms to identify all NE in toy community metabolic models that are analogous to classical games and predict reasonable interactions not captured by joint-FBA, OptCom and the dynamic method DMMM. We further validate NECom by analyzing a previously reported algae-yeast co-culture system, which shares a typical crossfeeding feature of lichen: fungi providing CO_2_ for photobionts, which fix nitrogen for fungi. NECom is able to predict important trends observed from the experimental data with minimal input from the data and to provide insights into the metabolic states of the two organisms.

**Figure 1.**
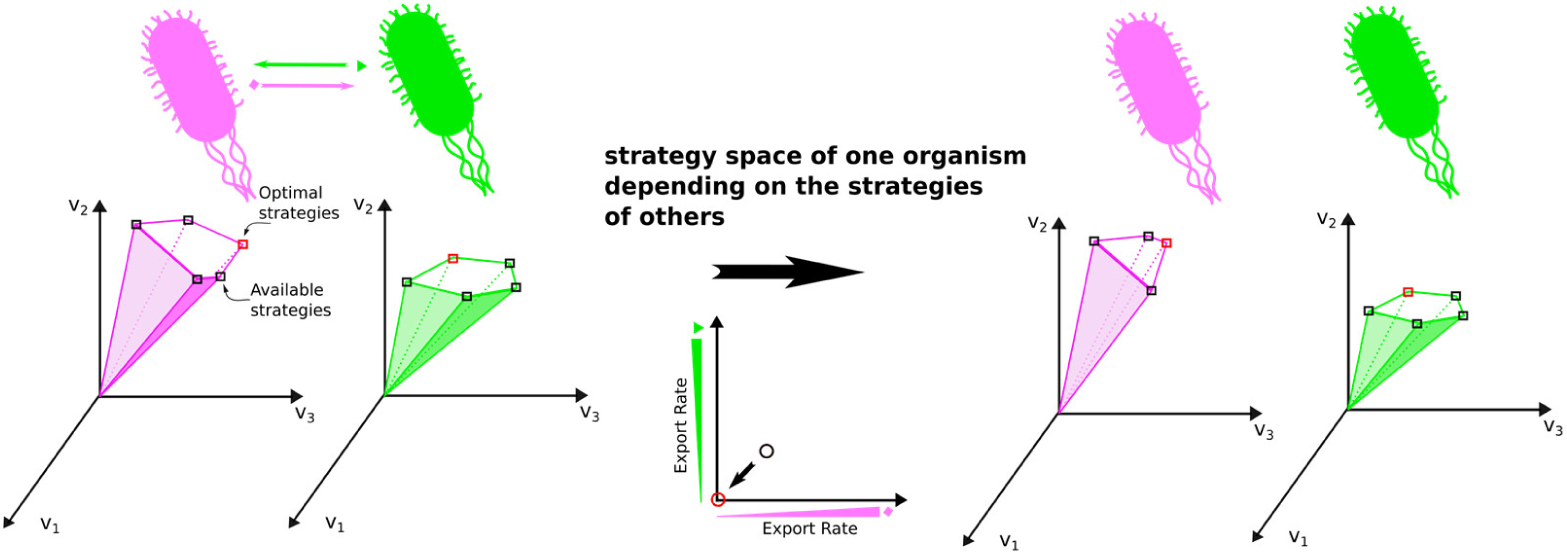
Schematic diagram of interactive strategy space of two community members. An available strategy for an organism is any feasible flux distribution satisfying the pseudo-steady-state condition and reaction directionality constrained by the availability of extracellular metabolites, which can be affected by other community members. Interactive strategy space causes the complexity of microbial metabolic games, distinguishing microbial metabolic game from most traditional games.

## RESULTS AND DISCUSSION

### A bi-level optimization approach for predicting global Nash Equilibrium of metabolic interactions

A bi-level optimization framework termed <u>N</u>ash <u>e</u>quilibrium predictor for microbial <u>com</u>munity (NECom) was developed. The biggest advance of NECom (Figure *2*) in comparison with previous methods is the absence of ‘forced altruism’ constraint. While the inner problem is essentially the Flux Balance Analysis (FBA) problem (48), the primary innovation lies in modeling how community metabolites available to the member microbes depend on the exchange profile (strategy) of each microbe. We proved mathematically that NECom prediction can guarantee Nash equilibria for its community members (**Supplementary Note**). With community biomass production as the outer-level objective function, solutions consistent with ESS or strict Nash equilibria can be found (in some cases ESSes depend on the actual competition of substrates which in turn depends on uptake kinetics that is not modeled in this study). The community-level objective function was used in all NECom predictions in this study. The derivation of NECom is presented in **MATERIALS AND METHODS**

**Figure 2.**
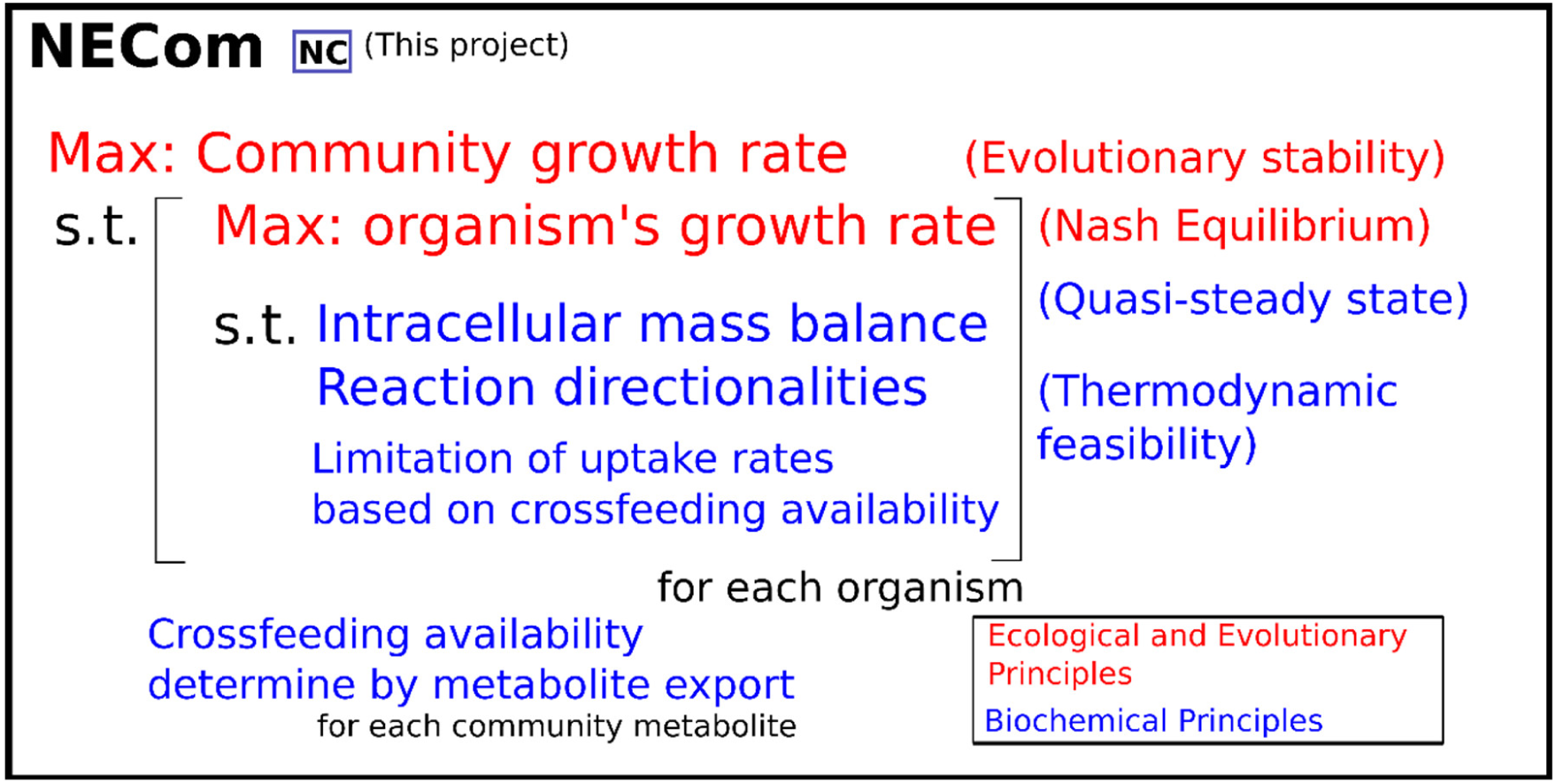
Schematic diagram of the NECom formulation.

### Predictions by NECom consistent with classical games analysis

#### Sub-optimal growth in a community of prototrophs (the prisoner’s dilemma)

To test and compare the proposed algorithm with other algorithms in a simple and clear way, four toy models including classical cases in game theory were analyzed. Case 1 is a community model consisting of two individual species (or mutants) with equal abundance named sp1 and sp2 (Figure 3a). Each species can yield ATP through converting substrate S to intermediate P. ATP is subsequently required to convert P to biomass precursors A and B, which can be freely exchanged between the two species. The difference between them is that sp1 produces B more efficiently (consuming 1 ATP versus 3 ATP for sp2), while sp2 produces A more efficiently (consuming 1 ATP versus 3 ATP for sp1), reflecting the existence of pathways with various yields that are commonly observed in nature. Intuitively the optimal strategy for the community is that each species cross-feeds each other with the metabolites produced at higher efficiency, i.e., sp1 supplies B to sp2, while sp2 feeds A to sp1, coinciding with the prediction by joint-FBA and OptCom (the lower flux map in Figure 3a). However, it is not a NE since either species, say sp1, can have a mutant that does not excrete B but keeps consuming A and increases its own biomass production. Using NECom, we predicted that the two species will grow at a lower rate and exchange nothing (the upper flux map in Figure 3a). Since there is no way for each species to grow faster given no export by another species, this is a NE. The dynamic method DMMM predicted the same results. In analogy to classical matrix game, a payoff matrix corresponding to different cross-feeding states (Figure 3a) was constructed. The payoff matrix indicates that maximum growth without cross-feeding is a strict NE (the ‘[0, 0]’ cell on the top left of the payoff matrix in Figure 3a; see also the explanation on the top right of Figure 3). A strict NE by the Maynard Smith’s first condition is also an evolutionary stable strategy (ESS), which is a NE not invadable by other mutants (or strategies) (49). Any other strategies combinations including the complementary cross-feedings (the [4.17, 4.17] cell in Figure 3a) predicted by Joint-FBA/OptCom are not stable and will eventually evolve to the NE. The interaction in the toy community model analyzed above is characterized as a game called the ‘prisoner’s dilemma’, because the NE is all members choosing ‘defect’ instead of ‘cooperate’, which NECom and DMMM correctly predict. The ‘forced altruism’ implied by the formulation is the cause for the mutualistic prediction by joint-FBA and OptCom because the formulation artificially prioritizes the metabolite export by each member for its partner over the optimization of its fitness in order to achieve higher community-level fitness.

**Figure 3.**
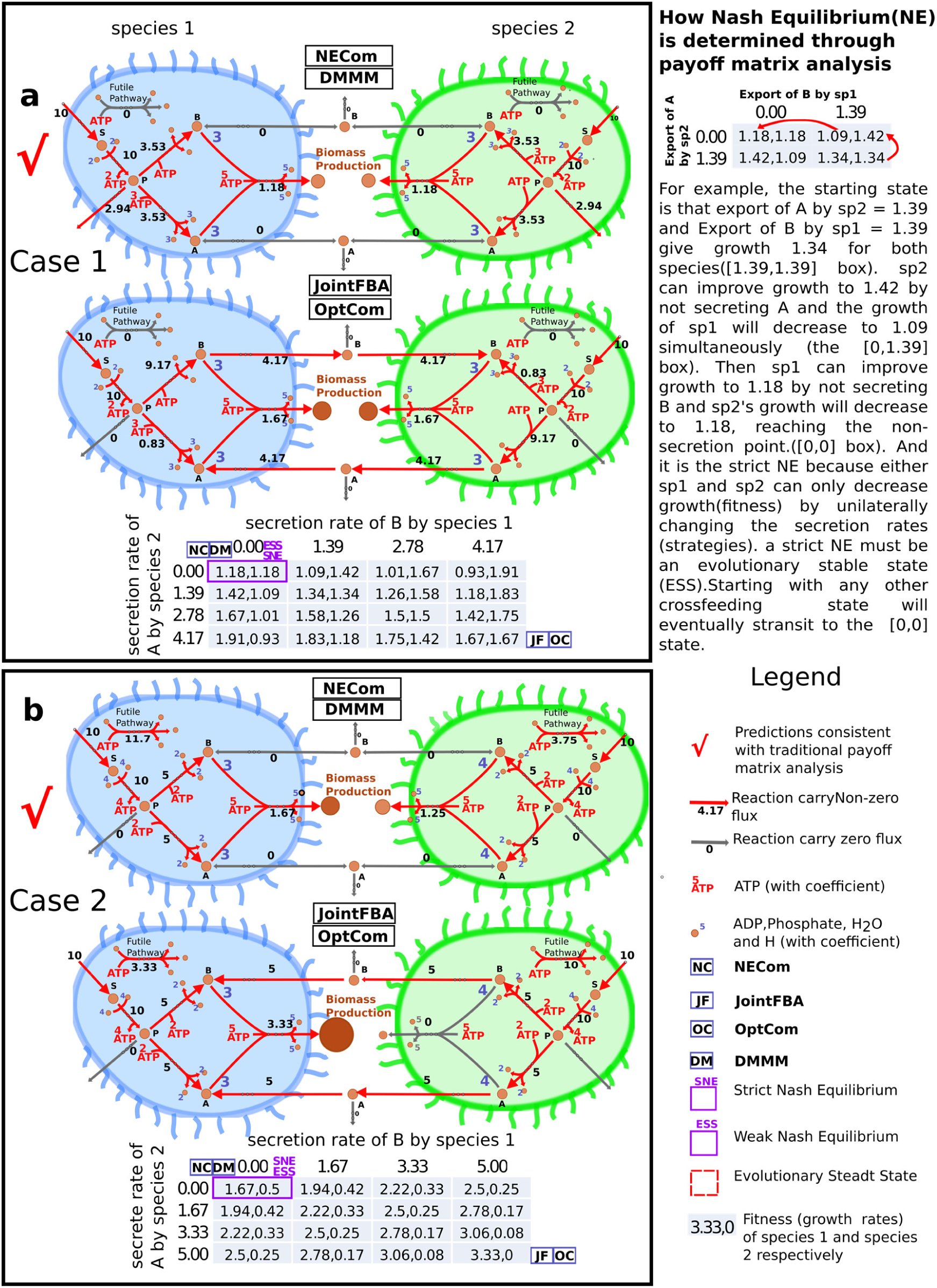
Flux distribution for toy models 1 and 2 predicted by different algorithms. The corresponding flux distributions predicted by Joint-FBA, OptCom, DMMM and NECom (NC) respectively. Growth rates of species 1 (sp1) and species 2 (sp2) calculated with FBA are shown in each cell in the payoff matrices for each case. **(a)** Toy community model 1 consisting of two members, both capable of producing biomass precursors A and B, but at different ATP costs. **(b)** Toy community model 2 consisting of two members, both capable of producing biomass precursors A and B, but has different biomass yields on these two precursors. Only NECom and DMMM can predict the ESS derived from the traditional payoff matrix approach in these two cases where the ESS is not mutualistic.

#### Differential biomass yields between members

Case 2 is a slight variant of case 1: the two organisms have different biomass yields on the same precursors. This reflects a phenomenon common in nature that different species have the same biomass precursors such as essential amino acidsbut the biomass yields on them are different. NECom indicates that non-cross-feeding is the strict Nash equilibrium/ESS (Figure 3b upper). However, Joint-FBA and OptCom predict that sp2 exports all of its biomass precursors A and B to support the growth of sp1 (Figure 3b lower). Such results are the consequence of formulations that invoke ‘forced altruism’, which determines that doing so achieves the highest overall growth, despite that sp2 needs to completely sacrifice its own fitness.

#### Evolutionarily stable mutualism between auxotrophs

Auxotrophy is common in microbial communities. Unlike the previous cases, auxotrophs rely on complementary cross-feeding for growth while the cross-feeding metabolites such as amino acids can be costly to produce. In such case how does cooperation become favorable or would defect still be the outcome despite no growth? Can cheaters and donors co-exist? To test how different algorithms predict this scenario, in case 3, we used the same setup as in case 1 but removed the reaction for synthesizing metabolite A in sp1 and the reaction for synthesizing B in sp2 (Figure 4). This mutualist-cheater case is a classical question of interest in ecology (50-53). The payoff matrix in Figure 4 shows that all strategy combinations, including mutualism, commensalisms, and non-crossfeeding are NE, and Satisfying exactly the Maynard Smith’s second condition (54), the mutualism (cell [5, 5]) is the ESS but there is no strict NE (see the explanation on the top right of Figure 4). Therefore from the view of evolutionary game theory, the non-producing cheaters in this case can still co-exist with producers although not being able to outgrow them. We checked that all NE identified by the payoff matrix are feasible solutions of NECom. The ESS was identical to the prediction of NECom, OptCom and Joint-FBA. However, DMMM predicts no growth and no crossfeedings. This false prediction is caused by the algorithm design, which is basically performing FBA for each organisms in each time-step. Each organism is only indirectly connected via metabolites in the medium and has no way to ‘know’ potentially more mutually favorable strategies from other organisms, which evolution might select for. This example also explains at the level of metabolic resource why mutualism or commensalism is possible even if it costs members some resource: because when a particular metabolite is not growth limiting, exporting the metabolite does not penalize the organism’s fitness (e.g., sp1 is limited by A so exporting B does not penalizing sp1’s growth and increases sp2’s growth at the same time). We applied NECom and DMMM to all N pairs of *E. coli* mutants auxotrophic for different amino acids that were experimentally tested in (55) and verified the same trend of predictions by each of the two methods (data not shown). DMMM does not predict any E. coli mutant exporting the amino acid essential to its partner organism, even when the amino acid it is auxotrophic for is available in the initial media. The predicted growth rate for each member becomes zero after all amino acids initially present in the media are depleted even if there is remaining glucose. In contrast, NECom predicts mutualistic cross feeding between the auxotrophs.

**Figure 4.**
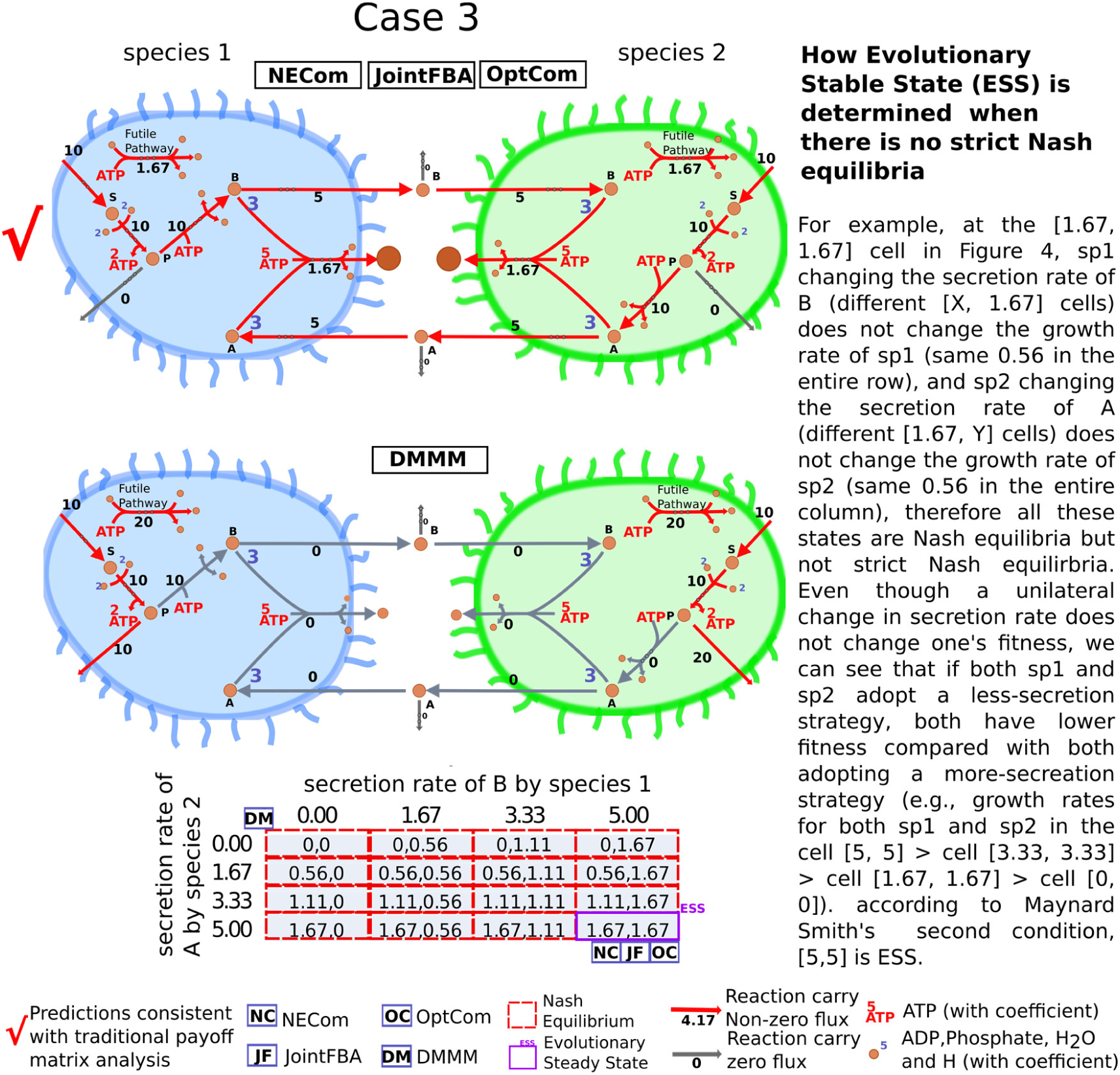
Flux distribution for toy model 3 predicted by different algorithms. Toy community model 3 consists of two members, one capable of producing biomass precursors A but auxotrophic for B, the other capable of producing biomass precursors B but auxotrophic for A. Only NECom, OptCom and Joint-FBA are able to predict the ESS derived from the traditional payoff matrix approach in this case where the ESS is mutualistic.

#### Co-existence of mutualists and cheaters (snowdrift game)

Then we tested the algorithms applied to a community with more than two members in case 4 (Figure 5a). The community was constructed by adding sp3, an exact copy of sp2, to the toy model in case 3. sp2 and sp3 are dependent on the export of B by sp1 and sp1 is dependent on the export of A by the other two members. An interesting question is whether the symmetric pair sp2 and sp3 will both produce A at same time. Nash equilibrium predicted by NECom indicates that sp2 and sp3 will always choose opposite strategies: when sp2 produces A (cooperation), sp3 does not (cheating) and vice versa (note the symmetry between sp2 and sp3 in Figure 5b). This defining feature categorizes the interactions as a typical ‘snowdrift game’ between sp2 and sp3. The prediction explicitly demonstrates the conclusion of our analysis of case 3 that mutualists and cheaters can co-exist in a NE. This case also suggests that NECom has the potential to predict species/strains level heterogeneity caused by evolution.

**Figure 5.**
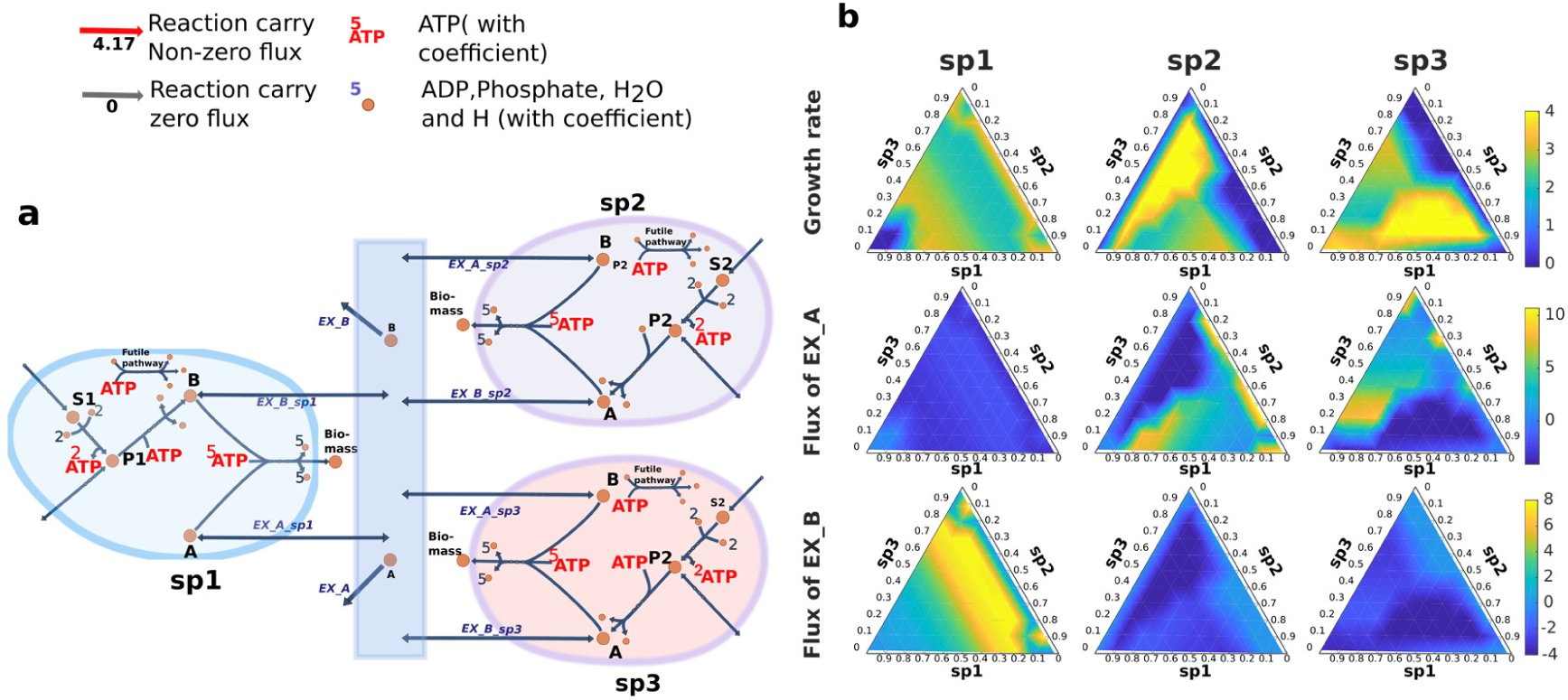
Toy community model 4. **(a)** Toy community consisting of three organisms. sp1 and sp2 are the same as in Toy model 3 and sp3 is a copy of sp2. sp2 and sp3 both depend on B produced by sp1, and sp1 depends on A produced by either or both of sp2 and sp3. **(b)** Growth rates and the cross-feeding reaction rates for each species under different relative abundance distributions. Note the symmetry in the column of ternary plots for sp1 and the antisymmetry between the columns for sp2 and sp3.

### Analysis of co-culture of *C. reinhardtii* and *S. cerevisiae* using NECom

Through toy model analysis NECom was demonstrated to be able to predict results consistent with traditional game-theoretical analysis. Next NECom was applied to a realistic algae-yeast co-culture system. Hom and Murray reported that two phylogenetically distant species *C. reinhardtii* and *S. cerevisiae* can establish obligatory symbiosis in simple media (56). Their data showed that *C. reinhardtii* almost always had a higher abundance than *S. cerevisiae* in all 80 experiments with different initial substrate concentrations. Within a wide range of initial concentrations of two key substrates (nitrite and glucose), these two species can co-exist for an extended time period (≥10 days, Figure 6). To elucidate how nutrient availability and the underlying metabolic interactions between the two organisms shaped the community growth pattern and the implication on the system stability, we performed a series of analyses by applying NECom (Figure 6) to a community model integrating the genome-scale metabolic models for *S. cerevisiae* (Yeast 7.6) (57) and *C. reinhardtii* (*i*Cre1355) (58). First we show the predictive capability of NECom for this system. Then from the simulation results we provide insights into growth-limiting nutrients and cross feeding metabolites and how they contribute to the predicted negative frequency dependent dynamics which stabilizes the community.

**Figure 6.**
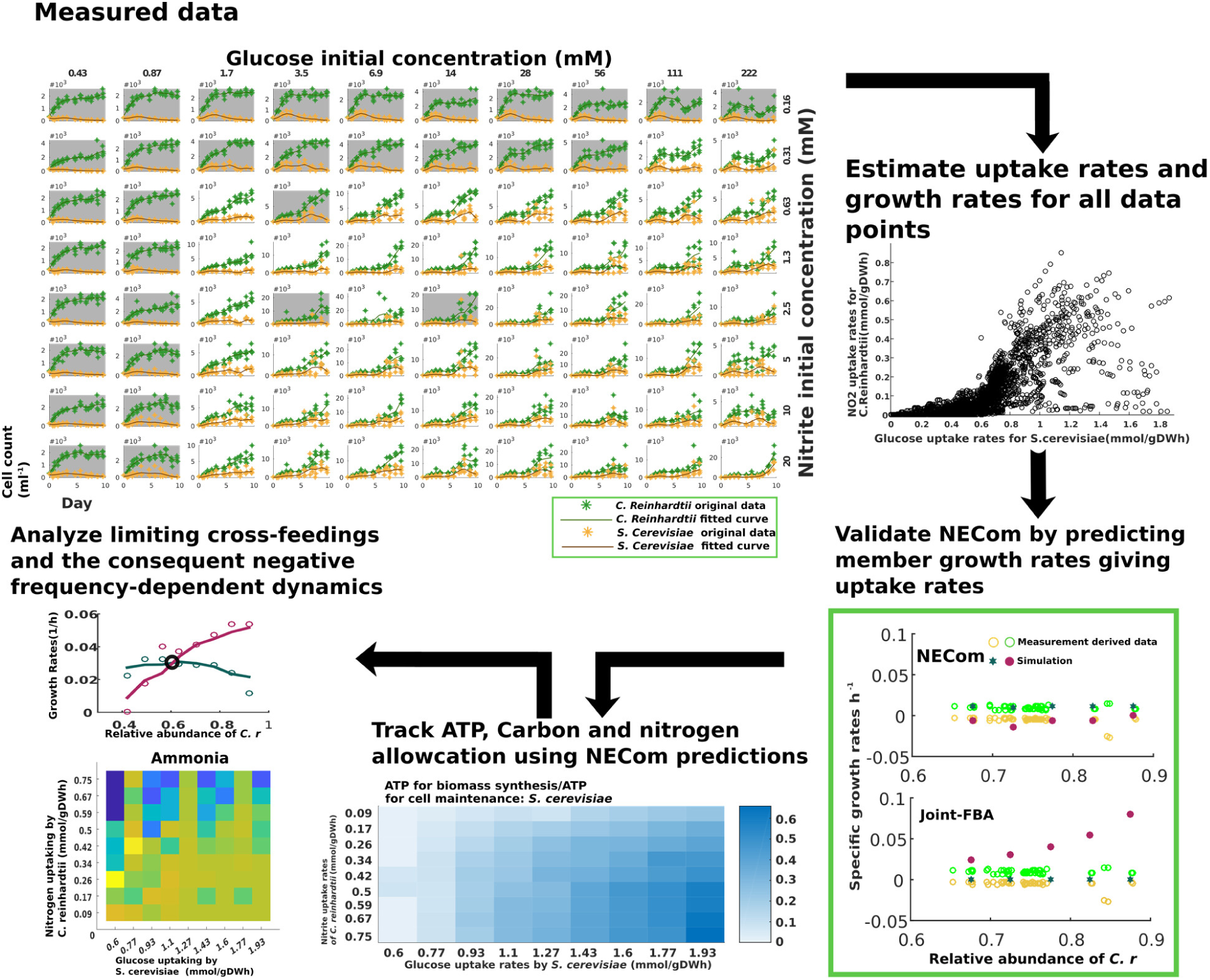
Workflow of analysis of co-culture of *C. reinhardtii* and *S. cerevisiae* using NECom.

#### Improved growth prediction by NECom

In order to validate the applicability of the NECom method to the *C. reinhardtii*-*S. cerevisiae* system, growth rates given uptake rates derived from experimental measurements (see **Supplementary Note**) were predicted using both NECom and Joint-FBA under a wide range of conditions (Figure 7a, b). The average root mean squared error (RMSE) for NECom is 3.1e-3 (*h*^-1^) for predicting the growth of *C. reinhardtii* and 8.5e-3 for the growth of *S. cerevisiae* respectively, achieving 63.5% and 81.7% reduction when compared to those predicted by Joint-FBA (statistical significance in reduction verified by Z-test, n =63, p-value = 3.12e-7 and 1.56e-23 respectively). The growth rates of *S. cerevisiae* predicted by Joint-FBA are generally significantly higher than the experimental results, leading to *S. cerevisiae* predicted as a generally faster grower while the experimental results indicate the opposite in majority of the cases (see Figure *7*b). When the glucose uptake of *S. cerevisiae* is low (<0.6 mmol/gDWh), Joint-FBA predicts positive values instead of the experimental negative values. These errors are caused by ‘forced altruistic’ constraints, which force *C. reinhardtii* model using photons to convert low energy carbon containing metabolites (primarily CO_2_) released by *S. cerevisiae* to various energy richer substrates subsequently to be consumed by *S. cerevisiae* (see the example in Figure *7*c). We confirmed this by maximizing *S. cerevisiae*’s growth and *C. reinhardtii*’s growth individually under the same condition using Joint-FBA and found *S. cerevisiae*’s maximum growth rate 2.7 times greater than *C. reinhardtii*’s. Therefore, ‘forced altruistic constraints’ unreasonably weaken the role of individual fitness and lead to inaccurate predictions. For the prediction by NECom, although in most cases it agrees well with the measurement-derived data, in some cases (e.g., nitrite uptake = 0.225 mmol/gDWh, glucose uptake = 0.725 mmol/gDWh) *S. cerevisiae*’s growth is underestimated while *C. reinhardtii*’s growth is overestimated. One possible reason is over-estimated uptake rates of CO_2_ by *C. reinhardtii*. When nitrogen is in excess, a higher CO2 uptake causes more-than-actual ammonia converted to biomass and less-than-actual ammonia exported for *S. cerevisiae*’s growth. The discrepancies are likely due to the absence of uptake kinetics in the current study.

**Figure 7.**
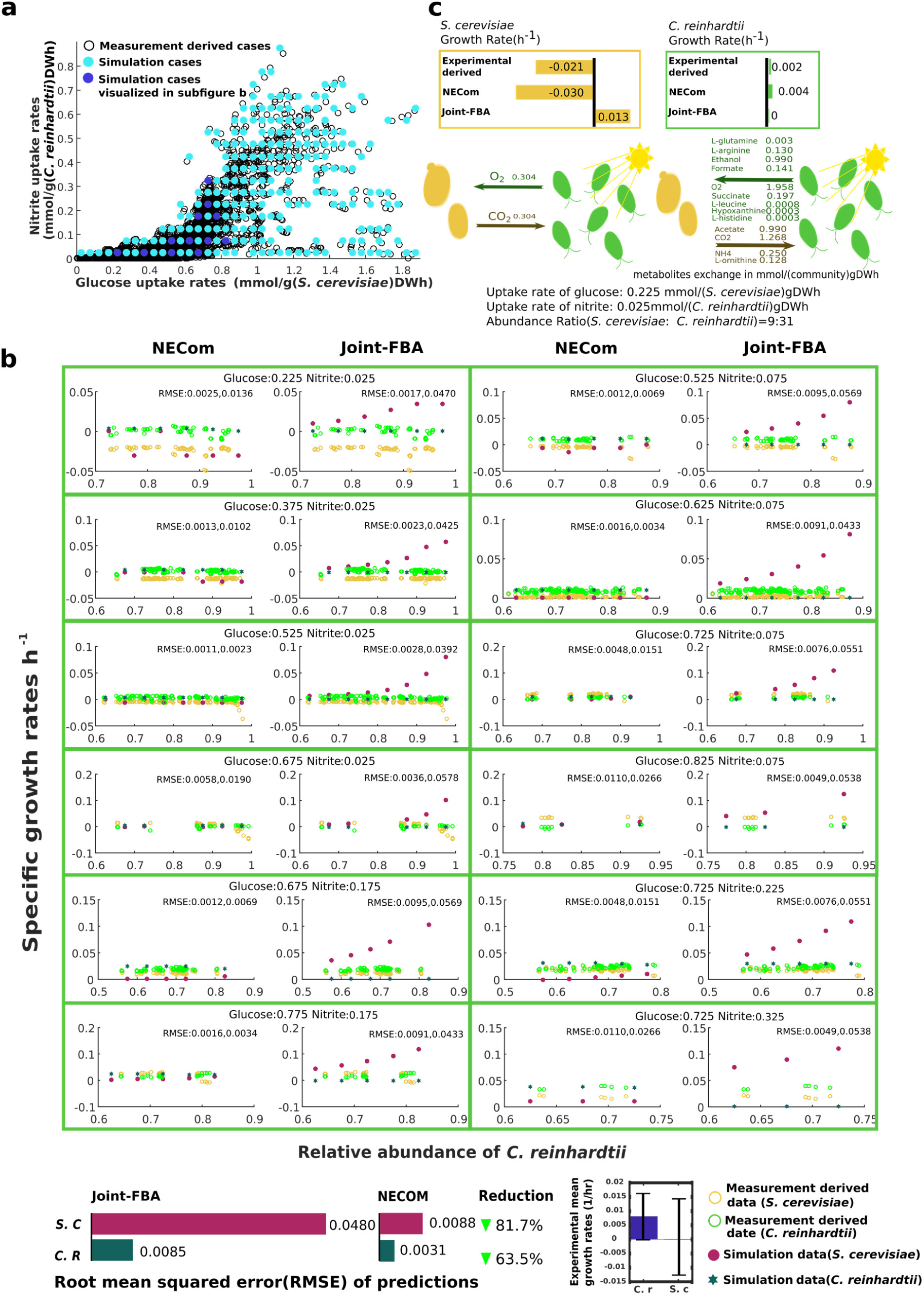
Verification of NECom and comparison with Joint-FBA. **(a)** Distribution of experimental and predicted cases in terms of the substrate uptake rates. Hallow circles represent the 3221 experimental cases (see Supplementary Note, noting that an axis of abundance is not presented in the figure, one circle can represent multiple cases with different abundances). Glucose and nitrite uptake rates were estimated based on the elemental composition of biomass and time series of cell counts (see Supplementary Note). (Dark and light) blue filled circles represent 488 simulation cases, which represent the experimental cases and with representativeness scores assigned (see Supplementary Note). **(b)** Simulation results of example cases (dark blue filled circles in (a)) using NECom and Joint-FBA compared with experimental cases. Rooted squared mean error (RSME) are calculated in each case. Weighted average RSMEs for both simulation methods are shown at the bottom. **(c)** Example demonstrating the difference of prediction between NECom and Joint-FBA in terms of predicted cross-feeding metabolites and its associated fluxes. RSMEs are labeled.

#### Impacts of substrate uptake rates on the relative growth between members

To the contrary of our impression that *S. cerevisiae* usually grows faster, experimental data (see Figure *8*c) suggests that in 76.7% of the cases, *C. reinhardtii* is the faster grower in this co-culture and usually has positive growth while *S. cerevisiae* could have negative growth especially when its glucose uptake rate is low(see Figure *7*b). The lower the glucose uptake is, the more the advantage of *C. reinhardtii* can have. Though at high glucose uptake rates (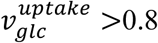 mmol/gDWh, see Figure *8*c), *S. cerevisiae* grows faster. We suspected that the algae being able to catch up and surpass the yeast at the depletion of glucose is important to the stabilization of the community and likely controlled by the change of substrate uptake rates. NECom predictions capturing the aforementioned phenomena (Figure *8*d) were then used to analyze the underlying mechanism.

**Figure 8.**
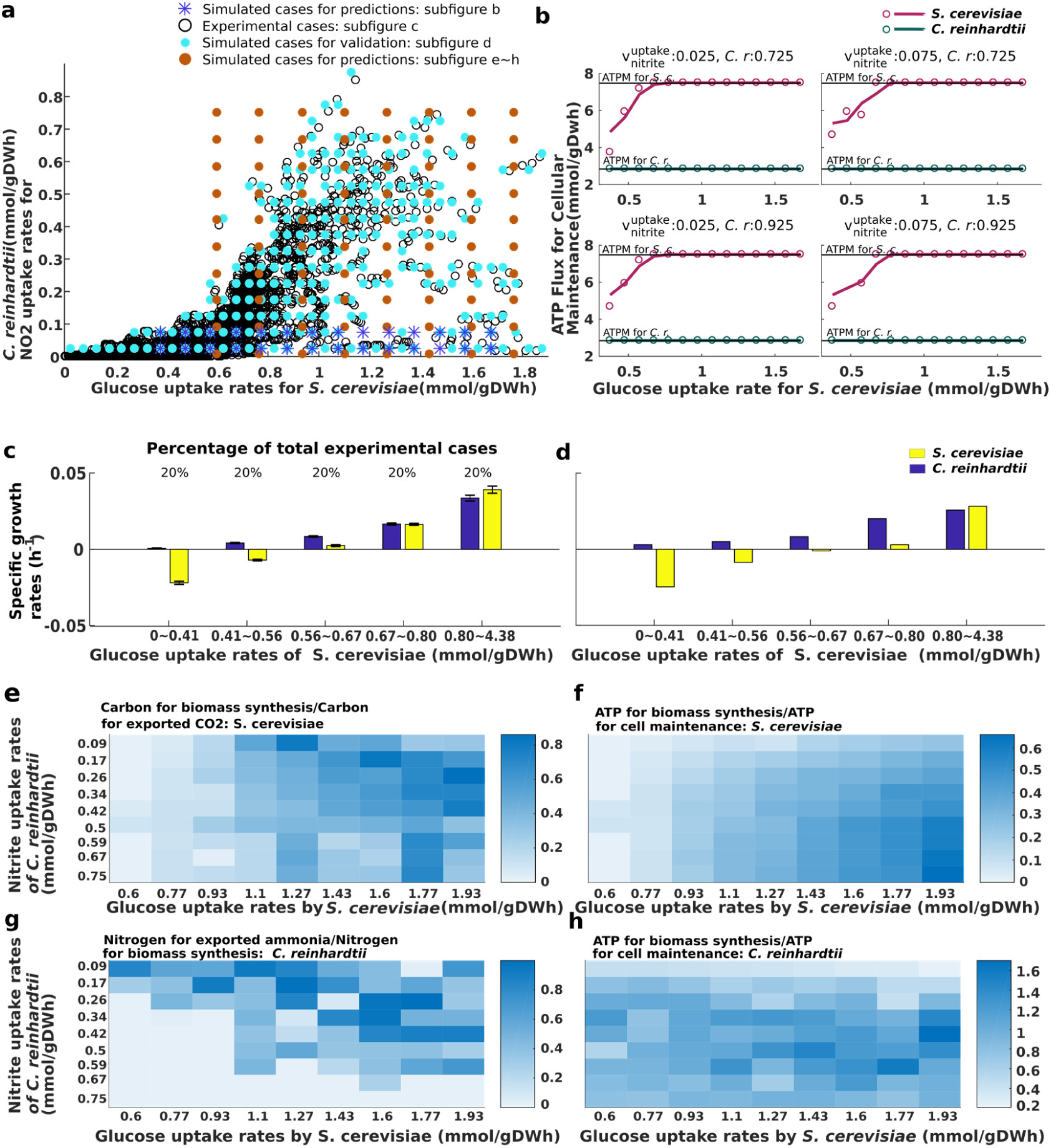
Mass and energy flow analysis. **(a)** Experimental and simulation data distribution. Hollow circles and light blue filled circles are set up in the same way as Figure 7a. Brown filled circles represent NECom predictions at various *C. reinhardtii*’s relative abundances, 631 cases in total (see **Supplementary Note)**. Dark blue asterisks represent simulation cases for ATP analysis shown in (b). For each point, simulations were performed at two different *C. reinhardtii* abundances (0.725 and 0.925). **(b)** ATP consumed for the cellular maintenance of *C. reinhardtii* and *S. cerevisiae* as a function of the glucose uptake rate for *S. cerevisiae* with the nitrite uptake rate and abundance of *C. reinhardtii* fixed at different values. **(c)** Average specific growth rate of *C. reinhardtii* and *S. cerevisiae* from experimental data (represented by the hallow circles, which are grouped into five intervals of glucose uptake rate, each interval includes 20% of the experimental data. **(d)** Weighted average specific growth rate of *C. reinhardtii* and *S. cerevisiae* over simulations represented by light blue filled circles (488 cases in total). The weight is the representativeness score explained in **Supplementary Note. (e - h)** NECom-predicted distribution of elemental flows and energy flows, using simulation cases (represented by brown filled circles in **a**) covering a wide range of glucose and nitrite uptake rates where experimental cases are sparsely distributed.

Using NECom predictons, we examined the always-positive growth of *C. reinhardtii*. Since the range of the experimental glucose uptake rate is wide enough for analysis only at low nitrite uptake rates, two representative low nitrite uptake rates, 0.025 and 0.075 mmol/gDWh, were selected for further analysis (67% of the experimental cases has estimated nitrite uptake rates < 0.1 mmol/gDWh). The maximum flux through the ATP maintenance (ATPM) reaction for each species was computed. If the ATPM flux is lower than a minimal value, which is the non-growth-associated maintenance (NGAM) value (2.85 mmol/gDWh for *C. reinhardtii* and 7.5 mmol/gDWh for *S. cerevisiae*), the growth rate will be negative. We found that the minimal ATPM flux is always satisfied for *C. reinhardtii*. However, for *S. cerevisiae*, when the glucose uptake rate drops below 0.8 mmol/gDWh (Figure *8*b), the ATPM flux starts to drop below the minimal requirement, the gap keeps increasing as the glucose uptake rate continues to decrease. One probable reason for this difference lies in the different source of energy used for fueling ATPM by the two species. For *C. reinhardtii* the energy source is photons whereas for *S. cerevisiae* the energy originates from metabolizing glucose. In the experiment, illumination was constant but the glucose concentration had different initial values and decreased over time, supporting our analysis.

Then, we investigated the experimental trend that *C. reinhardtii* narrowed its growth rate gap with *S. cerevisiae* and then surpassed it with the decrease of glucose uptake caused by the depletion (see Figure 9). The carbon flux to the biomass of *S. cerevisiae* and nitrogen flow to the biomass of *C. reinhardtii* have opposite trends. Based on NECom prediction, in response to the decreasing glucose uptake rates *S. cerevisiae* has a decreasing ratio of ATP for biomass synthesis to the less variant ATP for cellular maintenance, leading to a decreasing growth rate (Figure *8*f). In the meantime, The ratio of carbon flow to *S. cerevisiae* biomass over that to exported CO_2_ decreases and is responsible for more carbon available for *C. reinhardtii* to synthesize biomass (Figure *8*e). This is consistent with the following limiting substrate analysis, where CO_2_ is revealed to be a limiting substrate to *C. reinhardtii* (see Figure 9d).

**Figure 9.**
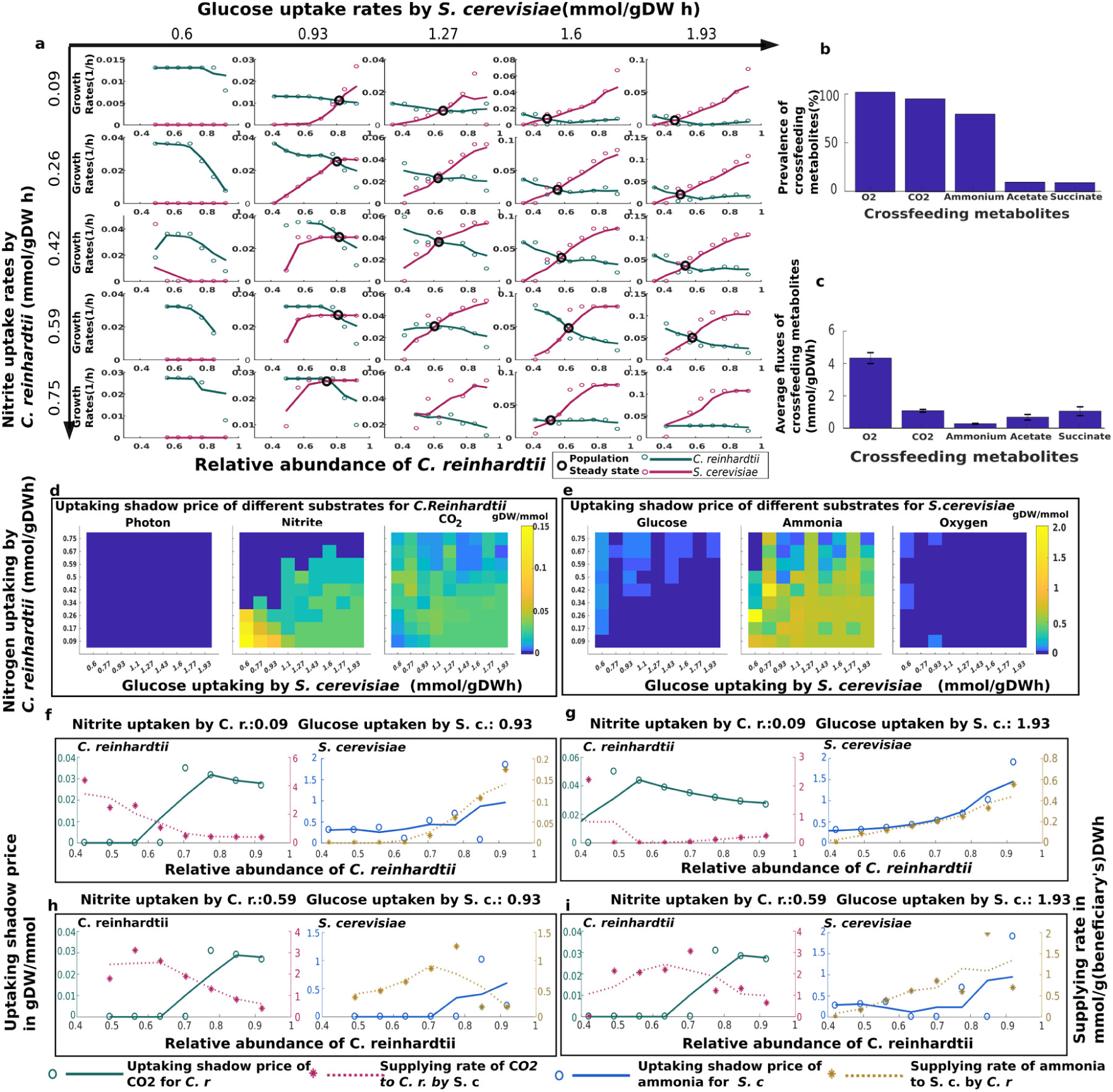
Shadow price based limiting substrate analysis. Simulation cases referred in this figure are those represented by brown filled circles in Figure 8a. **(a)** Growth rates of *C. reinhardtii* and *S. cerevisiae* as functions of the relative abundance of *C. reinhardtii*, at medium to high glucose uptake rates. **(b - c)** Prevalence and average flux of each cross-feeding metabolites across all simulation cases. **(d - e)** Average shadow price of major substrate uptake (including cross-feeding metabolites) by *C. reinhardtii* and *S. cerevisiae* under various nitrite and glucose uptake rates. **(f - g)** Shadow prices and supply rates of cross-feeding CO_2_ for *C. reinhardtii*, and ammonia for *S. cerevisiae* as functions of the relative abundance of *C. reinhardtii*.

#### Predicted negative frequency-dependent dynamics caused by limiting cross-feeding metabolites

The analysis thus far shows the ability of NECom to capture the growth trend of the members and provides some metabolic insights into the potential reason for the observed trend. A general question of interest for microbial communities is whether the community is stable and how the environmental conditions impact the stability. To complement the limitation that the measured data were concentrated in a small range of low substrate uptake rates, we extrapolated the predictions by NECom over a large range of relative abundance of *S. cerevisiae* and *C. reinhardtii* to address this question. NECom predicts that with higher glucose uptake rates (>0.75 mmol/gDWh) the growth rates of the two species will have opposite trends to the relative abundance of *C. reinhardtii* and often there exists a point of equal growth rate (black circles in Figure 9a), where the faster and slower growing species are swapped. Such a growth-abundance relationship is negative frequency-dependent, which stabilizes the co-existence of interacting species by dynamically evolving to the equal-growth-rate population steady state. An interesting observation is that the population steady state shifts toward the left on the abundance axis with the increase of glucose uptake rate for *C. reinhardtii* (Figure 9a), indicating that the growth of *S. cerevisiae* gains advantage over *C. reinhardtii* with an increasing glucose uptake rate. The prediction is consistent with the experimental data (Figure *8*c).

We speculated that the changing availability of some key substrates might be the reason for the negative frequency dependent dynamics and the shift of population steady state. To test our speculation, all cross-feeding metabolites and externally sourced substrates need to be considered. We asked which metabolites were the major substrates used by the two species. Among the simulation cases at various glucose uptake rates, nitrite uptake rates and abundances (represented by brown dots in Figure *8*a), cross-feeding metabolites observed include O_2_, CO_2_, ammonia, acetate and succinate. The former three metabolites occur in 93% of the total cases (Figure 9b). Therefore, O_2_, CO_2_, ammonia together with external sourced substrates including photon, glucose and nitrite were selected for further analysis. Since only limiting substrates can have impact on community members’ growth, in order to specify which substrates are growth-limiting, shadow prices for the uptake of each metabolites were extracted from NECom solutions (see Supplementary Note). A positive shadow price indicates the corresponding metabolite has impact on cellular growth at a given state and the value indicates the intensity of such impact. We found that for *C. reinhardtii*, CO_2_ is a limiting metabolite under almost all simulated conditions (Figure 9d), nitrite have significant limiting effect when both nitrite and glucose uptakes are low. Photon is generally not limiting. For *S. cerevisiae*, only ammonia exhibits significant limiting effect (Figure 9e).

We then focused on analyzing how CO_2_ and ammonia affect the growth rates of this system. Figure 9f-i indicate that with increase in the abundance of *C. reinhardtii*, CO_2_ supplied by *S. cerevisiae* decreases, causing decrease in the growth rate of *C. reinhardtii* since CO_2_ is the limiting substrate confirmed by the positive shadow price curve. In the meantime, increase in ammonia supply from *C. reinhardtii* contributes to the increasing growth rates of *S. cerevisiae*, since ammonia uptake has increasingly positive shadow prices. Therefore, the opposite trends in the supply rate of each limiting substrates are responsible for the observed negative frequency-dependent dynamics. Shadow price analysis can also explain why the population steady state changes with the glucose uptake rate. Figure 9a shows that with the increase in glucose uptake rate, the two growth-abundance curves becomes steeper at low *C. reinhardtii*’s abundance (<0.6mmol/gDWh), causing the cross point shifting to the left. The steepness of a growth-abundance curve is proportional to the shadow price for limiting substrate uptake. Comparison of Figure 9g and Figure 9f reveals that, fixing the nitrite uptake rate at 0.09 mmol/gDWh and increasing the glucose uptake rate from 0.93 mmol/gDWh to 1.93 mmol/gDWh result in an increase in shadow price for CO_2_ uptake by *C. reinhardtii* from 0 to 0.02-0.04 gDW/mmol when the abundance of *C. reinhardtii* is below 0.6. Similarly Figure 9i and Figure 9h indicate that if the nitrite uptake rate is fixed at 0.59 mmol/gDWh, shadow prices for ammonia uptake by *S. cerevisiae* will increase from 0 to 0.25 gDW/mmol when the abundance of *C. reinhardtii* is below 0.7.

### Relations to previous methods

A game-theoretical method has been proposed recently to determine Nash equilibria for interacting organisms using metabolic models(43). In that proposed method, the Black Queen hypothesis (59) was tested by checking whether the existence of auxotrophic mutants is favored by evolution if there is leakage of metabolites. Metabolite leakage was predefined and uptake of leaky metabolites was determined using a distribution rule. FBA was then performed for each genotype in the population to compute the payoff matrix which was subsequently used for Nash equilibrium determination as well as dynamic simulation using the replicator equation. While the results elegantly captured *S. cerevisiae*’s invertase system and suggested potential auxotrophic pairs that can stably co-exist, similar to other dynamic methods, performing FBA for each individual genotype, or more generally organism, does not allow individuals to ‘sense and response’ to each other via changes in the metabolic environment. Therefore, the possible interactions observed are confined to the cross-feeding of the predefined leaky metabolites. In the current study, the Black Queen hypothesis is not invoked (though it can be). Also, by clearly defining the concept of microbial metabolic games and the unique strategy space which is a continuous flux space constrained by the strategies of other players, NECom is able to search the inter-dependent continuous flux space to determine Nash equilibria among all available interactions by using a bi-level framework, in which each organism optimizes its own fitness, under the substrate uptake constraints shaped by the strategies of other community members. In relation to previous algorithms for simulating community metabolic models, OptCom (35) also employs a bi-level optimization framework but the main innovation of NECom lies in the introduction of outer variables to constrain the uptake only for the inner problems instead of both uptake and export (see Methods for the mathematical details). SteadyCom (38) implements a population steady state but the single objective function also unavoidably imposes forced altruism on the community the same way as joint-FBA does. We have not implemented the population steady state in the formulation but it can still be analyzed by fixing the relative abundances at different values as performed in the algae-yeast case study. In Table 1 some key features of NECom and current algorithms are compared.

**Table 1.**
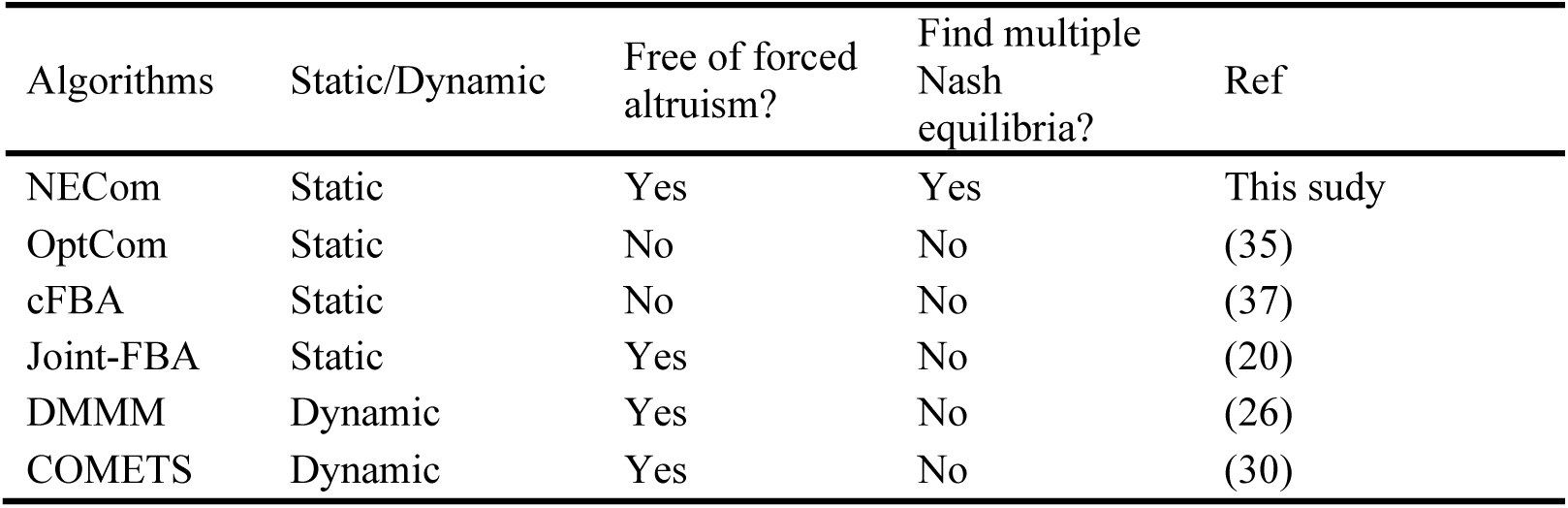
Comparison of representative algorithms for predicting microbial metabolic interactions

| Algorithms | Static/Dynamic | Free of forced altruism? | Find multiple Nash equilibria? | Ref |
| --- | --- | --- | --- | --- |
| NECom | Static | Yes | Yes | This study |
| OptCom | Static | No | No | (35) |
| cFBA | Static | No | No | (37) |
| Joint-FBA | Static | Yes | No | (20) |
| DMMM | Dynamic | Yes | No | (26) |
| COMETS | Dynamic | Yes | No | (30) |

In conclusion, we developed a bi-level non-convex mixed integer optimization algorithm called ‘NECom’ to predict the Nash equilibrium of multiple species interacting at the metabolic level. NECom was shown to predict Nash equilibria of different types of classical games correctly in toy models that previous methods’ predictions cannot guarantee. The main reason for this difference is that NECom does not contains any ‘forced altruism’ that computationally forces individual species to fulfill other species’ metabolic need before their own. As proof of concepts, NECom was used to analysis a well-documented co-culturing system of *C. reinhardtii* and *S. cerevisiae*. With minimal information from the experimental data, NECom is able to reveal that the decreasing glucose concentration vs. the constant photon availability provides significant growth advantage of *C. reinhardtii* over *S. cerevisiae*; CO_2_ and ammonia are growth-limiting cross-feeding substrates for *C. reinhardtii* and *S. cerevisiae* respectively; and the negative frequency-dependent growth pattern is caused by opposite trends of the supply rates of each species’ limiting substrates.

## MATERIALS AND METHODS

### Derivation of NECom

Let *N* be the set of organisms in a microbial community. For each organism *n* ∈ *N*, let *I*_*n*_ be the set of metabolites, *J*_*n*_ be the set of reactions, 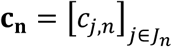 be the objective coefficient vector, and *v*_*j,n*_ be the flux of reaction *j* ∈ *J*_*n*_. The FBA problem for organism *n* can be formulated as:

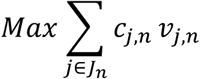

subject to:

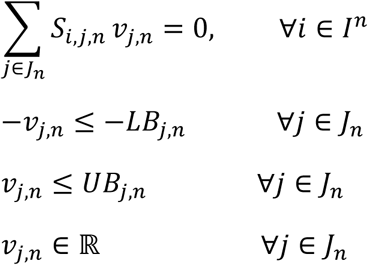

where 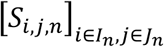 is the stoichiometric matrix, *LB*_*j*_ and *UB*_*j*_ are the lower and upper bounds for the flux of reaction *j* respectively. The equality constraint represents metabolite balances at pseudo-steady state. The inequalities include bound constraints due to reaction directionality, substrate availability and the requirement of non-growth-associated-maintenance (NGAM) modeled as a minimum flux through an ATP hydrolysis reaction, which is usually called ATP maintenance (ATPM) in metabolic models.

The FBA problem will be the inner problem of NECom for optimizing the fitness of individual organisms. A set of outer-level variables to indicate the community metabolites available for uptake are then introduced and connected to the exchange flux variables of individual organisms. Let 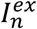 be the set of extracellular metabolites in the community, 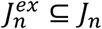 be the set of exchange reactions between the community and organism *n*. Define an index mapping function 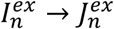 such that *com*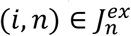 map extracellular metabolite 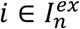 to its corresponding exchange reaction. For each individual organism, we split the exchange reaction flux for a possible cross-feeding metabolite *v*_*com*(*i,n*),*n*_ into two non-negative continuous variables: the uptake rate 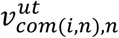 and the export rate 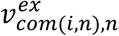:

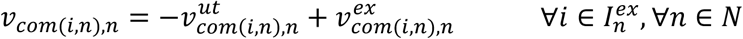

To explicitly model the mutual dependency in terms of inter-organism metabolite exchange, uptake of a metabolite by an organism is possible only if there is surplus of the metabolite in the medium after the consumption/production by the rest of the community:

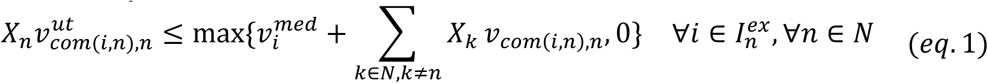

where *X*_*n*_ is the relative abundance of organism *n* (a pre-set parameter), which is multiplied by specific uptake rate 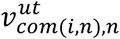 to correctly normalize the exchange with the community, and 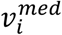 is the maximum community uptake of metabolite i from the medium. Here to avoid forced altruism, 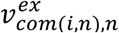 is designated as an inner variable independent of the outer problem, while the uptake rate 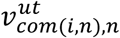 is also an inner variable but depends on the net availability of metabolite i in the community. In this way we can ensure that individual organisms have ‘autonomy’ over their metabolite exports. Maximization of the community fitness in the outer level cannot force an organism to produce a certain metabolite unless FBA determines that the metabolite is necessary for, or at least not undermining the maximum growth of the organism. This modeling constraint is novel to our best knowledge. The max function in the *eq*.*1* is linearized by introducing an outer continuous variable *β*_*i,n*_ and an outer binary variable *δ*_*i,n*_:

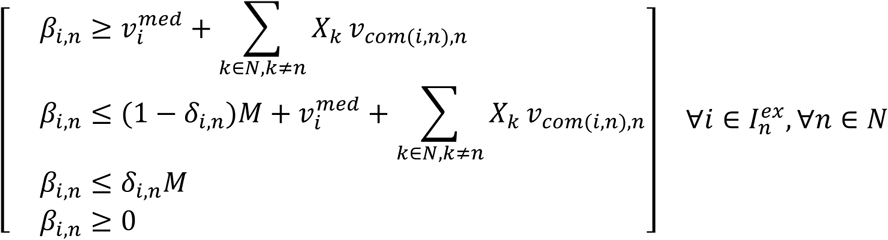

Where *M* is a large constant. Combining the inner problem and outer problem the complete formulation of NECom can be written as:

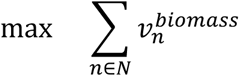

subject to

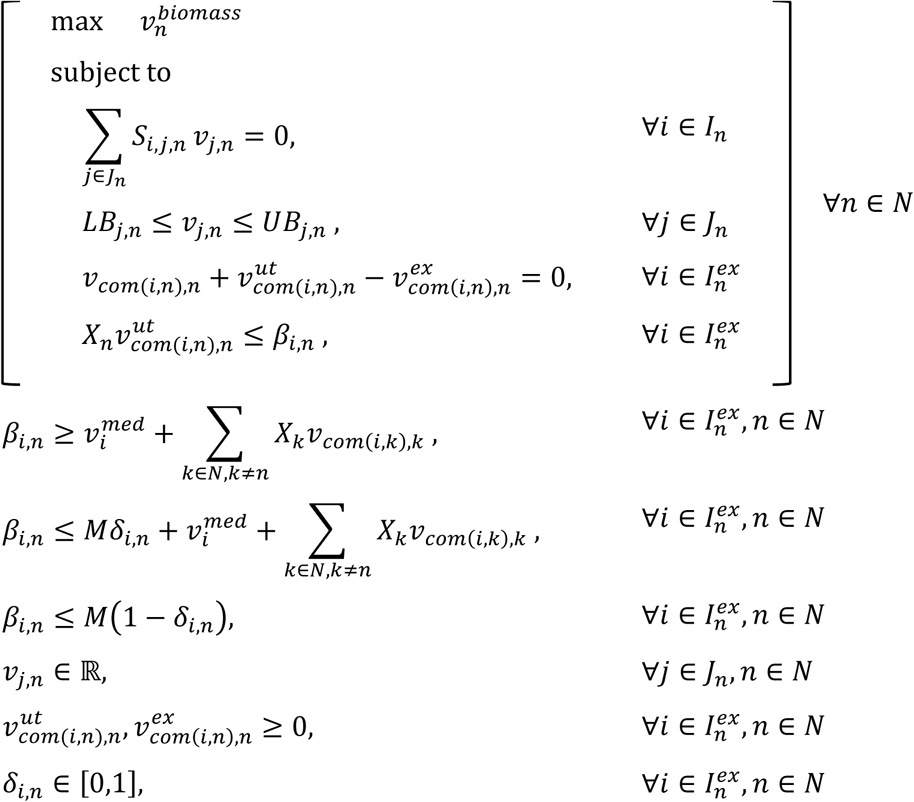

The only situation where the above algebraic system is not feasible is that one or more members cannot satisfy the minimal NGAM, which is represented as a direct ATP hydrolysis reaction with a pre-defined lower bound in model. Biologically this suggests the occurrence of cell death. In order to enable NECom to predict this phenomenon, we separate community members into two sets based on NECom solution: let *N*^*G*^ be the set of community members which have adequate substrates for uptake to satisfy their NGAM, *N*^*D*^ be the set of the rest of the community members that are dying due to insufficient nutrients. For the latter case, the default FBA problem is infeasible. Instead of maximizing biomass production, we relax the NGAM constraint and maximize the ATP hydrolysis activity that an organism can generate to minimize cell death, resulting in NECom enabling negative growth prediction (NEComNG) as:

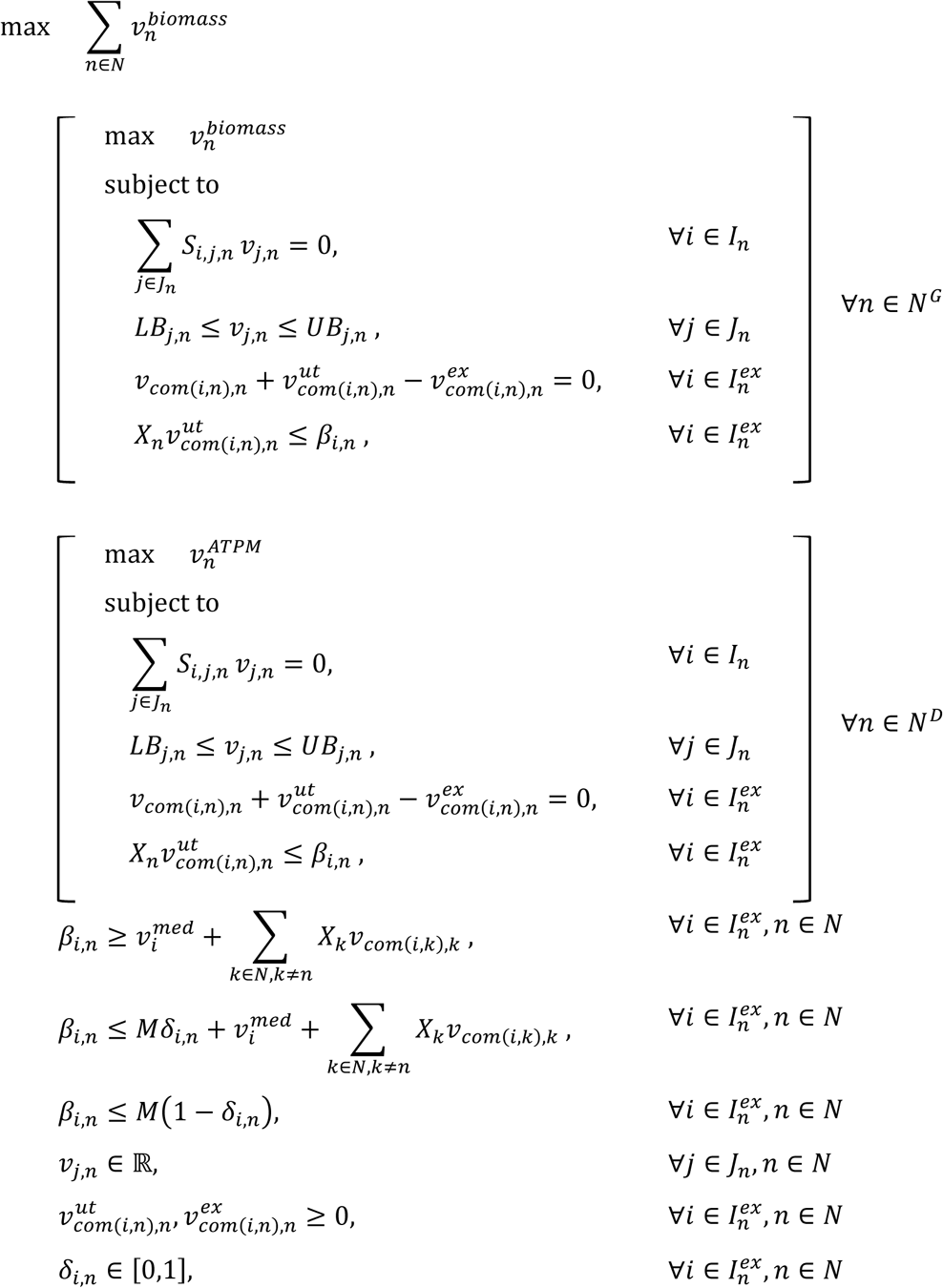

The transformation of the bilevel optimization problem into a mixed integer non-linear problem is detailed in **Supplementary Note**.

### Software used

Baron (60) was employed to solve all non-linear and mixed integer non-linear optimization problems. CPLEX and GLPK were used to solve linear problems. Pathway maps were originally generated in Escher Map (61). All scripts were written in Matlab.

### Code and data availability

Data files and example codes will be available on https://github.com/Jingyi-Cai/NECom.git

## Acknowledgments

We acknowledge Dr. Wenyan Yuan, Dr. Erik F. Y. Hom, Dr. Guang Cheng, Dr. Xueying Ni and Ying Zhu for their useful comments.

## Author contributions

Conceptualization, J.C and S.H.J.C; Methodology, J.C and S.H.J.C; Software, J.C; Analysis and Investigation, J.C and S.H.J.C; Writing – Original Draft, J.C and S.H.J.C; Writing – Review & Editing, S.H.J.C, T.T, and J.C; Visualization, J.C and S.H.J.C; Supervision, T.T and S.H.J.C.

## Competing interests

The authors declare no competing interests

## Notes

### Competing Interest Statement

The authors have declared no competing interest.

